# HazardPyMatch: A Tool for identifying reproductive and other hazards in scientific laboratories

**DOI:** 10.1101/2025.03.14.643365

**Authors:** Emily M. Parker, Anastasia-Maria Zavitsanou, Clara Liff, Danique Jeurissen, Nour El Houda Mimouni, Isabella Succi, Eric Rogers, Marianna Liistro

**Author notes:** These authors contributed equally (Emily M. Parker, Anastasia-Maria Zavitsanou).

## Abstract

Understanding and mitigating laboratory hazards is essential for fostering safe and inclusive research environments. However, conducting risk assessments can be challenging and time-consuming, especially for scientists who have new or specific concerns about hazard susceptibility, such as pregnant women. In response, using reproductive hazards as our primary example, we developed HazardPyMatch, a laboratory hazard screening tool designed to be implemented in laboratories across scientific disciplines to support efficient hazard management. HazardPyMatch is an accessible and user-friendly tool that enables scientists to quickly and easily systematically identify chemical hazards in laboratory chemical inventories and categorize these hazards in laboratory protocols.

**Motivation:** A publicly accessible and easy-to-use resource did not exist to identify reproductive hazards in neuroscience laboratories. In response we developed the python workflow HazardPyMatch to address the need for a data-driven tool to catalog hazardous materials in laboratory settings.

## Introduction

Conducting laboratory risk assessments can be incredibly time-consuming, given the hundreds of chemicals and substances typically stored in individual laboratories. While University Environmental Health and Safety (EH&S) departments create valuable resources for assessing laboratory hazards and risk assessment, existing hazard identification tools built by EH&S are not publicly accessible^1^. While EH&S departments offer consultations to assess specific risks in individual laboratories, it is typically not the case that these consultations can be performed in complete confidence. Further, individuals may actively choose to delay or not to disclose sensitive health information in the workplace. We developed HazardPyMatch in response to the need for consolidated and easily accessible resources to identify hazardous materials in laboratory settings. HazardPyMatch is a publicly available data-driven laboratory hazard screening tool that allows individuals to quickly and easily categorize laboratory chemicals that may pose increased risk in unique or specific circumstances.

With just a chemical inventory and, optionally, a collection of laboratory protocol documents, users of our python-based software quickly obtain critical information on hazard identification and classification. We use Chemical Abstracts Service (CAS) Numbers, identifiers for locating safety information, and leverage programmatic access to PubChem’s chemical annotations including the Globally Harmonized System of Classification and Labeling of Chemicals (GHS), an internationally standardized framework for classifying chemicals to identify and sort chemical inventories based on hazard category^2^.

Here, we present HazardPyMatch and the results from our assessment of reproductive hazards in neuroscience laboratories, to specifically address the need for quick data-driven hazard identification for pregnant women working in neuroscience. We describe HazardPyMatch inputs and outputs, details about the top ten chemical hazards identified in our primary assessment, and brief interpretations of our findings. To explore the broader applicability of our tool, we evaluate a second hazard classification. As Zebrafish is an increasingly common model organism in scientific research, we present our findings on aquatic life hazards in neuroscience laboratories. This application demonstrates generalizability of our tool and that it can be applied to help identify hazards to animals in laboratory settings. We also demonstrate the HazardPyMatch’s use in evaluating reproductive hazards in a chemical inventory from a de-identified chemistry laboratory at Columbia University. Our tool is not limited to these applications; scientists interested in carcinogenicity in laboratory settings or aspiration hazards, among others, could greatly benefit from utilizing HazardPyMatch.

Hazard identification is a critical step in laboratory hazard assessment and risk mitigation. We aim for HazardPyMatch to serve in this fundamental role, as a starting point for scientists to quickly and easily access chemical hazard information crucial for mitigating risk to human and animal health. HazardPyMatch outputs provide users with the information necessary to perform important next steps – to perform targeted literature searches to expand knowledge of specific laboratory hazard risks and, most importantly, to seek specific advice from experts in EH&S departments on risk mitigation to advance safety and increase awareness in scientific environments.

## Methodology & Results

### Obtaining and Uploading a Chemical Inventory

Users navigate to https://pubchem.ncbi.nlm.nih.gov/ghs/ to read about and select GHS H-codes based on hazard interest. GHS H-codes are standardized hazard statement codes consisting of “H” for hazard, followed by 3 numerical digits to indicate hazard characteristics: hazard class, category and signal word. In order to run the pipeline, each user obtains and uploads a chemical inventory or list to HazardPyMatch. ChemTracker has been widely adopted across academic institutions and is recommended for accessing the most up-to-date chemical inventories for individual laboratories^3^. We suggest uploading your chemical inventory to Google Drive and using the provided python code with Google’s IDE Colab, or similar. A ReadMe is supplied with the code to guide users through the workflow. Intermediate python users can download the code package and interact with it on a local machine. After inputting the correct file paths, the user is prompted to input GHS H-codes of interest. HazardPyMatch will load the chemical inventory (**Figure 1A**) and fill in any missing CAS Numbers. Any chemicals lacking CAS Numbers (the majority of which are proprietary reagents from laboratory supply companies) are excluded. Our first HazardPyMatch application was to identify reproductive hazards present in neuroscience laboratories based on common neuroscience protocols. When prompted we supplied “reproductive toxicity” and “germ cell mutagenicity” GHS H-codes: H360, H360D, H360FD, H360F, H360fD, H360Df, H361, H361f, H361d, H361fd, H362, H340, and H341 (**Table 1**). These are collectively referred to henceforth in the text as “reproductive hazards.” We obtained and loaded a Zuckerman Institute ChemTracker chemical inventory in collaboration with the Department of EH&S at Columbia University. This large chemical inventory merged the chemical inventories of 38 laboratories at the Zuckerman Institute totaling 3165 unique and non-unique chemical name entries. 60 entries were excluded due to lack of CAS Number. 3105 chemicals proceeded to the hazard identification and categorization steps.

**Table 1.** GHS H-codes related to reproductive toxicity hazards. The table includes classifications for chemicals that may damage fertility, the unborn child, or breast-fed children, as well as those suspected of causing such damage. Codes also encompass potential genetic defects.

| **GHS H-code** | **Hazard Statement** |
| --- | --- |
| H360 | May damage fertility or the unborn child. |
| H360D | May damage the unborn child. |
| H360FD | May damage fertility. May damage the unborn child. |
| H360F | May damage fertility. |
| H360Fd | May damage fertility. Suspected of damaging the unborn child. |
| H360Df | May damage the unborn child. Suspected of damaging fertility. |
| H361 | Suspected of damaging fertility or the unborn child. |
| H361f | Suspected of damaging fertility. |
| H261d | Suspected of damaging the unborn child. |
| H361fd | Suspected of damaging fertility. Suspected of damaging the unborn child. |
| H362 | May cause harm to breast-fed children. |
| H340 | May cause genetic effects. |
| H341 | Suspected of causing genetic effects. |

**Figure 1.**
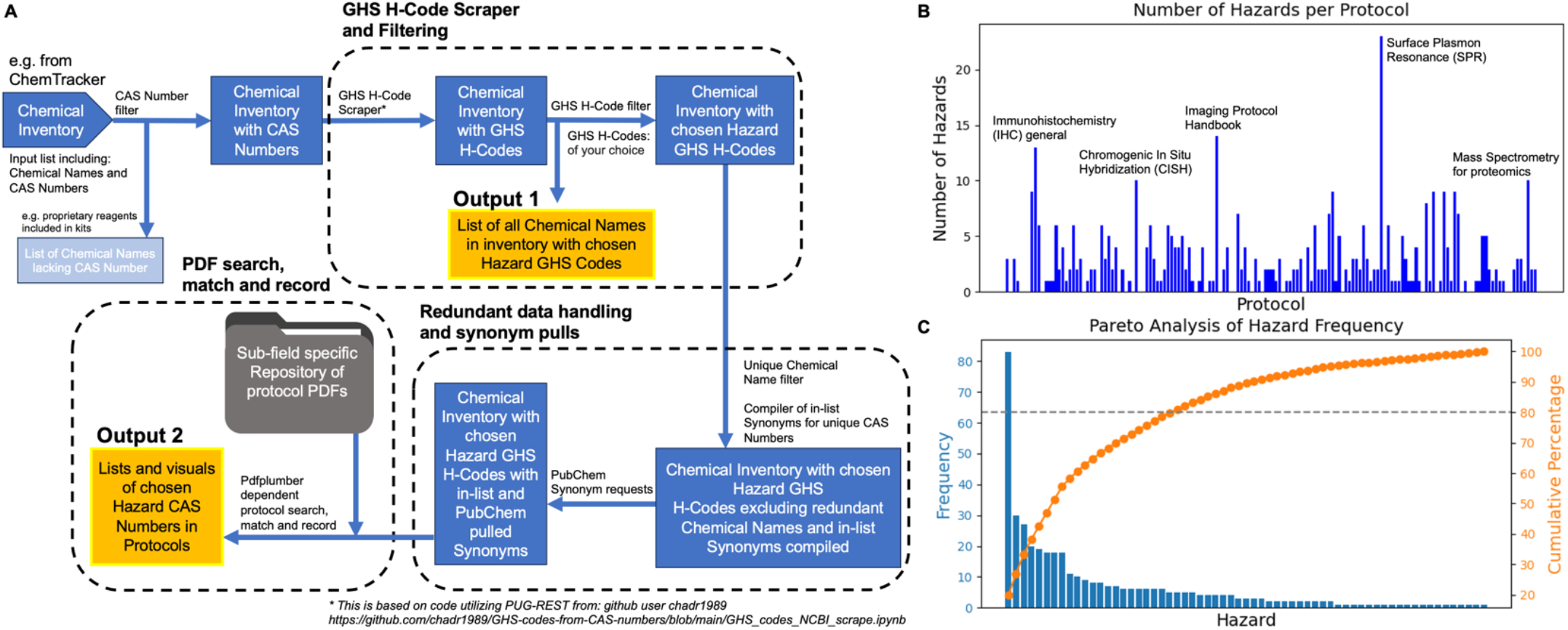
**HazardPyMatch is a python-based pipeline for extracting hazards in chemical inventories with output lists of hazards to be prioritized for risk mitigation and data visualizations.** **A)** A python-based software pipeline which takes a chemical inventory as input and extracts GHS H-codes from PubChem using CAS Numbers, filters chemicals by hazard classification of interest and reduces redundancy with synonym handling. It additionally searches protocol PDFs for hazardous chemicals and generates output lists and data visualizations which can then be further used for hazard risk assessment. **B)** Bar chart showing the Number of chemical hazards associated with each protocol. The distribution of the data generated from our primary application, reproductive hazards in neuroscience, highlights variations in hazard density with a range of 0-23 hazards in our repository of 152 common neuroscience protocols. **C)** Pareto chart displaying the frequency of individual chemical hazards (blue bars) along with the cumulative percentage of occurrences (orange line) indicating a small subset of hazards account for the majority of the occurrences, emphasizing key hazards (in **Table 2**) that should be prioritized for risk mitigation.

### Hazard Identification and Categorization

Next, chemicals are classified based on health and environmental hazard statements by integrating GHS H-codes with data from PubChem, one of the largest open-access chemical databases for chemical substances. This is achieved using the Power User Gateway Representation State Transfer (PUG-REST) service, a web interface for programmatically gathering information from PubChem^4,5^. The inventory is filtered to include only the chemicals associated with the user-defined H-codes and saved (**Figure 1A**, Output 1). GHS H-code filtering for reproductive hazard H-codes reduced our chemical inventory to 1085 unique and non-unique chemical names.

**Table 2.**
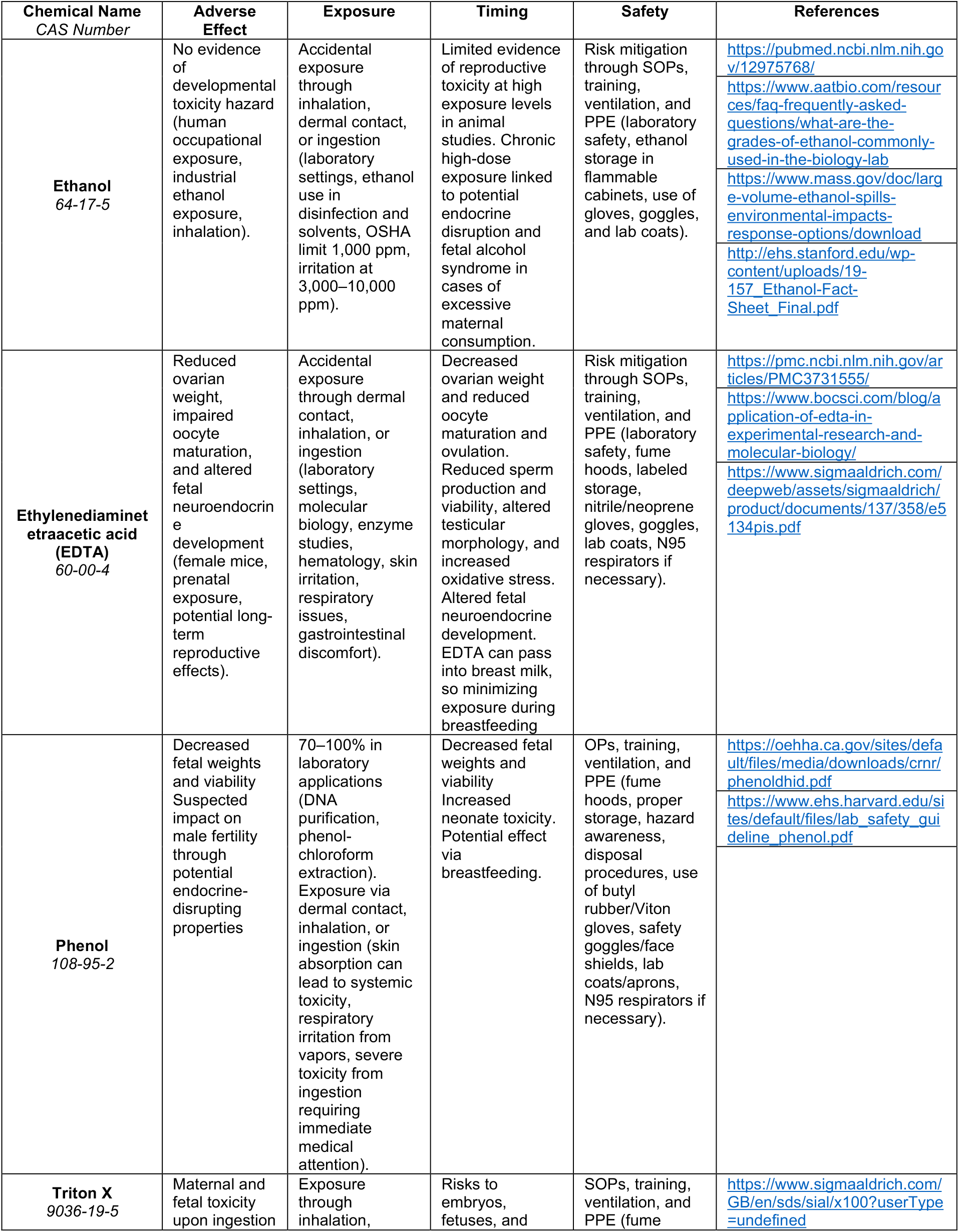
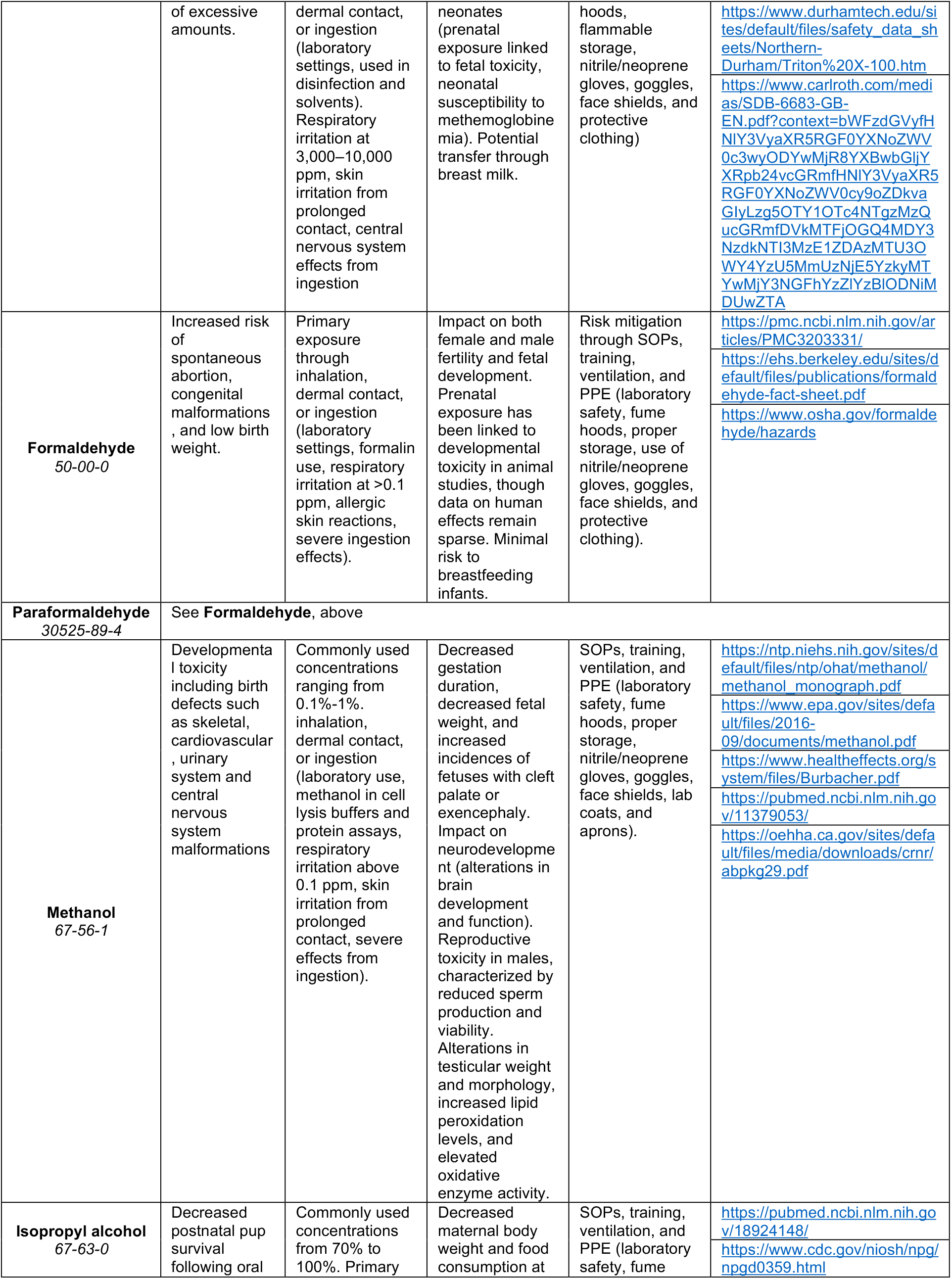
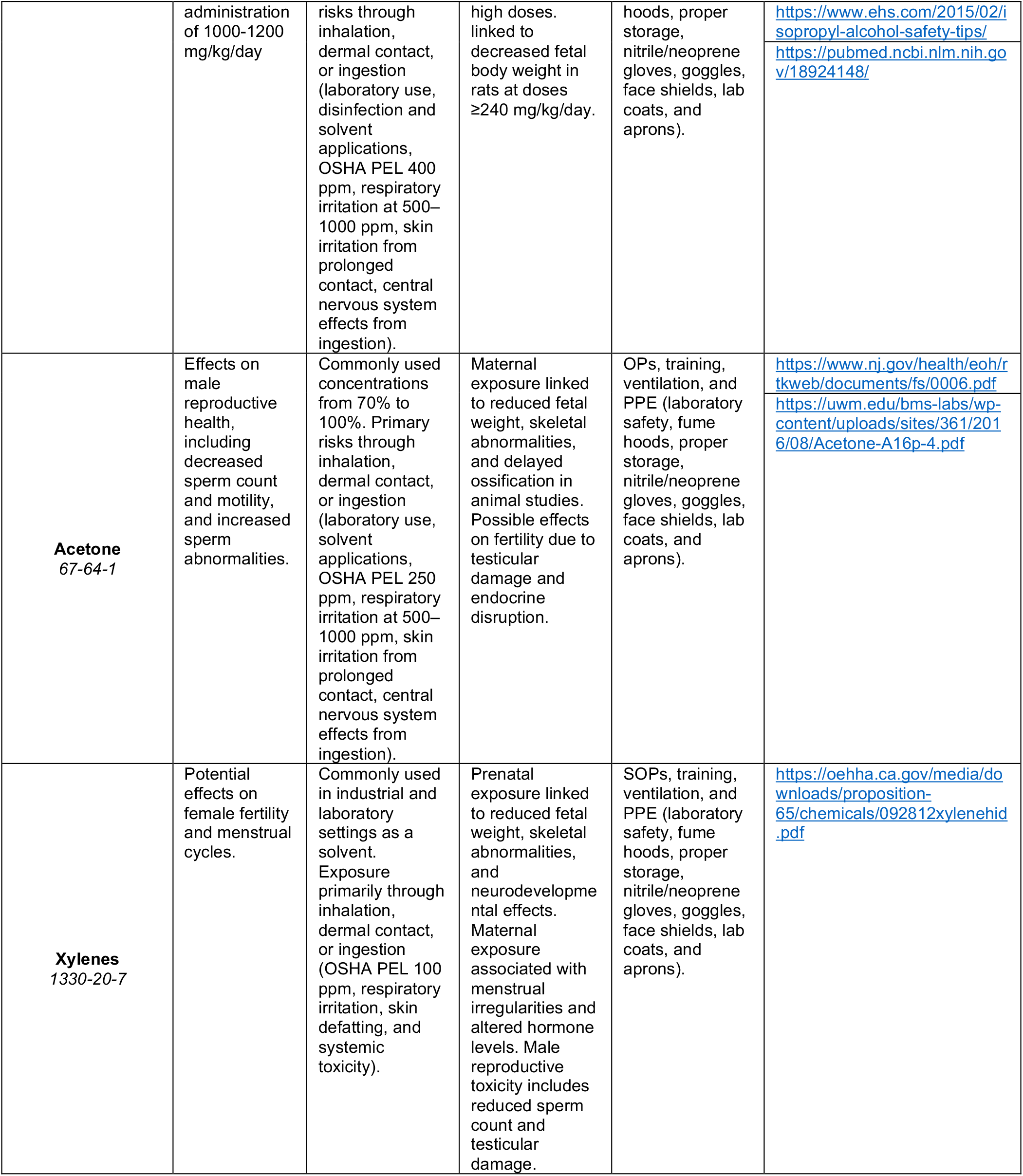
Systematic descriptions of the 10 most commonly used chemicals in neuroscience protocols, identified using GHS H-codes for reproductive toxicity. This table includes a literature review to summarize each chemical’s adverse effects, exposure, critical windows of susceptibility, and safety measures, with references supporting hazard assessment and risk mitigation strategies.

### Standardizing Chemical Nomenclature and Handling Redundancy

Each chemical inventory is then filtered down to unique chemical names, a redundant information handling step, and chemical name synonyms are compiled within the inventory itself. The PUG-REST service is utilized for a second time to integrate chemical name synonyms from PubChem into the inventory (**Figure 1A**), which is necessary given chemicals and substances are referred to by various and in some cases a vast number of aliases. These include but are not limited to trade name, abbreviation or International Union of Pure and Applied Chemistry (IUPAC) name. For our initial HazardPyMatch application on reproductive hazards in neuroscience laboratories, unique chemical name filtering and synonym integration reduced our chemical inventory to 144 unique entries.

### Mapping Chemicals to Experimental Protocols

Finally, users can optionally cross-reference the final chemical inventory to a repository of PDF protocol documents to catalog hazards within specific protocols. To do this, users upload protocol PDF files to a designated Google Drive folder, ensuring the folder path is correctly set. The PDFs, either from individual scientist’s own protocol collection or open-access platforms like protocols.io^6^, are analyzed using pdfplumber, a robust Python dependency for reading PDFs. This final step generates Output 2 from HazardPyMatch (**Figure 1A**), which includes a list of hazards in each protocol, a list of hazard terms with corresponding GHS H-codes, and exploratory data visualizations (**Figure 1B-C**).

We conducted a literature review to identify common experimental protocols in neuroscience and gathered feedback from scientists at the Zuckerman Institute of Columbia University to ensure comprehensive coverage of assays commonly used in our multidisciplinary scientific sub-field. We matched our hazards against 152 neuroscience protocols and identified 64 reproductive hazards that are commonly used in neuroscience research.

### Reproductive Hazards in Neuroscience Laboratories Output

We used our laboratory hazard screening tool HazardPyMatch to assess a large neuroscience institute-wide chemical inventory. Comparing Output 1 to our original ChemTracker input revealed that 34.5% of the chemicals listed in our chemical inventory from ChemTracker were linked to at least 1 of the 13 reproductive hazard GHS H-codes. Considering Output 2 we learned that the number of hazards in each neuroscience protocol ranged from 0 to 23, with 19% of the 152 neuroscience protocols we collected containing no identified reproductive hazards. This variation in hazard counts suggests that certain protocols may require increased risk mitigation. Further evaluating Output 2, we found that the top 10 neuroscience protocols with the highest total number of reproductive hazards comprised primarily biochemistry techniques (**Figure 1B**).

To better understand how frequently different hazards occur, we analyzed the distribution of hazards across all protocols from Output 2. Our analysis showed that a small number of specific hazards account for the majority of occurrences, meaning that certain hazards appear repeatedly across multiple protocols. For example, Ethanol was the most frequently occurring hazard in neuroscience protocols, appearing nearly 3 times as often as the next most common hazard. Ethanol’s role as a solvent, disinfectant, and dehydrating agent explains its widespread use in neuroscience protocols (**Figure 1C**).

We additionally provide **Table 2**, our systematic descriptions of the 10 most commonly used reproductive hazards in neuroscience protocols. Our results in **Table 2** integrate data from Output 2 from HazardPyMatch with additional information gathered through a literature review, detailing each chemical’s adverse effects, exposure, timing, and safety considerations. Relevant studies and information were identified through searches of NCBI, safety data sheets from reputable chemical vendors (e.g. Sigma-Aldrich) or universities and US or US state governmental institution websites (e.g. NIOSH and nj.gov). Users are responsible for supplementing information from literature searches to their own HazardPyMatch outputs.

### Extended Applications: Aquatic Life Hazards and Chemistry Laboratory Inventory

To demonstrate the versatility of HazardPyMatch, we used the Zuckerman Institute chemical inventory to evaluate a different GHS hazard category. We chose to evaluate aquatic life hazards to show how our tool could help assess risk management for laboratory animals, inspired by the growing use of Zebrafish as a valuable model organism in scientific research. We used the “toxic to aquatic life” GHS H-codes: H400, H401, H402, H410, H411, H412, and H413. After CAS Number filtering, aquatic life hazard GHS H-code filtering reduced our inventory to 511 unique and non-unique chemical names. The standardizing chemical nomenclature and redundancy handling step further reduced the inventory to 209 unique entries and protocol matching identified 37 unique aquatic life hazards found in the 152 neuroscience protocols we gathered. 19 chemicals are shared between the aquatic life hazards and reproductive hazard lists, a percentage overlap is 26.03%. A chi-square test was p<0.05, indicating the observed overlap is unlikely to have occurred by random chance.

Additionally, we demonstrate the applicability of HazardPyMatch to a different scientific discipline: chemistry. We utilized HazardPyMatch to identify reproductive hazards in a de-identified chemical inventory from a chemistry laboratory at Columbia University. The ChemTracker inventory was provided to us by Columbia University EH&S and contained 1510 total entries. 60 chemicals were excluded due to lack of CAS Number and 381 unique and non-unique chemicals remained after reproductive hazard GHS H-code filtering, 25.2% of the total chemicals in the ChemTracker version of the inventory. The redundancy handling step further reduced the inventory to a final list of 99 unique chemicals with 1 or more reproductive hazard GHS H-codes.

To assess the relevance of HazardPyMatch outputs, we compared our results to a comprehensive review of reproductive hazards in chemistry laboratories^7^. Prior research has identified commonly used organic solvents and heavy metals in these settings, many of which are known to be reproductive hazards. Notably, some organic solvents included in the aforementioned review lacked an associated reproductive hazard classification under GHS. In contrast, HazardPyMatch identified 8 organic solvents with reproductive hazard H-codes, highlighting potential updates or differences in classification criteria. Consistent with current GHS classifications, HazardPyMatch did not detect an association between Pentane and reproductive hazard classifications, whereas previous work had categorized it under GHS category 2 with no reported adverse effects. This discrepancy may be due to updates in GHS coding after the publication of earlier studies. Finally, HazardPyMatch did not produce false negatives within the scope of our analysis of the de-identified chemistry laboratory chemical inventory. Several chemicals identified as reproductive hazards in prior work were absent from the list we generated with HazardPyMatch, but this was consistent with their absence from the original ChemTracker inventory. These findings suggest HazardPyMatch effectively identifies reproductive hazards while successfully reflecting current GHS classifications.

### Limitations of Study & Discussion

Our efforts underscore the critical need for accessible and user-friendly resources that enhance safety in laboratory environments. Using reproductive hazards as an example, we developed HazardPyMatch, a laboratory hazard screening tool that can be adapted and implemented for any hazard type, across laboratories, institutions and scientific disciplines to support effective hazard management. This tool serves as a starting point for scientists, providing a foundation for informed decision-making while reinforcing the importance of consulting institutional safety personnel. We strongly encourage scientists to collaborate with their local EH&S teams to ensure compliance with safety protocols and maximize the effectiveness of hazard mitigation strategies.

By integrating chemical hazard data with experimental protocols, HazardPyMatch not only fosters safer research environments but also promotes inclusivity in STEM fields. HazardPyMatch was designed to empower scientists to make informed choices about personal health and the health of essential laboratory animal models. We developed this tool after noticing that pregnant scientists in neuroscience laboratories lacked a consolidated resource for identifying germ cell mutagenicity and reproductive toxicity hazards. We demonstrate the generalizability of our tool by evaluating hazards associated with aquatic life toxicity in neuroscience laboratories and then assessed the performance of our tool for evaluating germ cell mutagenicity and reproductive toxicity in a chemistry laboratory. Our tool is not limited to these applications. Other potentially interested parties include scientists interested in carcinogenicity in laboratory settings, aspiration hazards, or explosive hazards, to name a few. Furthermore, pharmaceuticals used in scientific research are generally excluded from chemical inventories, including ChemTracker, and we have not included them herein. The vast majority of drugs commonly used in academic research do have CAS Numbers thus HazardPyMatch can be used to for drug hazard cataloging and protocol matching. Positioned as a foundational resource, we aspire for this tool to contribute to a culture of safety and inclusivity in scientific research across all laboratory setting types.

While our tool provides a valuable resource for identifying laboratory hazards, it is important to recognize its limitations. First, HazardPyMatch’s scope is inherently limited to chemical hazard identification and protocol matching. This means that certain categories of substances, such as engineered nanomaterials and endocrine disruptors, which may lack CAS Numbers, fall outside its purview. Moreover, and most critically, while HazardPyMatch identifies hazardous chemicals, it does not account for crucial contextual factors such as exposure levels, routes of exposure, usage concentrations, and lab-specific safety measures, all of which significantly influence risk. Our findings on Ethanol perfectly exemplify this. Cross-referencing the Zuckerman Institute ChemTracker inventory, Ethanol was the most frequently occurring hazard in common neuroscience protocols using HazardPyMatch. The major reproductive risk Ethanol poses, however, is fetal alcohol spectrum disorder when Ethanol, as a component of a beverage, is chronically consumed by the pregnant mother^8^. Research has concluded that exposure to Ethanol, as an industrial chemical, is not associated with developmental toxicity^9^. Following hazard identification, contextual factors such as exposure levels, routes of exposure, usage concentrations, and lab-specific safety measures must be assessed manually, reinforcing the need for collaboration with EH&S experts to ensure proper risk mitigation.

HazardPyMatch integrates extensive synonym mapping utilizing PubChem synonyms to improve detection, however, it can identify false positives, where non-hazardous terms are mistakenly flagged. To address this, HazardPyMatch outputs include extracted text from source documents, enabling users to manually verify identified hazards. False positives are typically easily identifiable as common words or abbreviations, such as “Wood” appearing for Methanol’s CAS Number or “PEI” for Ethyleneimine. There also remains the possibility of false negatives—instances where hazardous chemicals are overlooked. Our testing identified one potential case, Isoflurane, which lacks associated reproductive hazard GHS H-codes. Although Isoflurane is not considered a reproductive hazard on Safety Data Sheets or under GHS, the National Institute for Occupational Safety and Health recommends Isoflurane levels are kept below the lowest detectable level. Additionally the UK provides a workplace daily limit of 50ppm, and reviews have not revealed adverse effects of Isoflurane when levels were below this limit^10^. Although studies have shown Isoflurane placental transfer, findings on potential reproductive and teratogenic effects including spontaneous abortion are inconclusive^8^. Nevertheless, users should remain cautious, particularly when working with infrequently utilized substances or those with less or inconclusive documentation.

In closing, HazardPyMatch serves as a valuable, publicly available resource for enhancing laboratory safety by providing scientists with critical insights into potential hazards. When integrated with targeted literature reviews and expert consultations, HazardPyMatch outputs empower scientists to take a data-driven proactive approach to hazard management. This not only helps mitigate risks but also fosters a culture of safety, responsibility, and inclusivity within scientific laboratory environments. By equipping researchers with the tools to identify and address hazards effectively, HazardPyMatch supports the creation of safer and more responsible scientific workplaces.

## Acknowledgements

We sincerely appreciate the support of the members of Zuckerman Institute Gender Inclusion Group for their guidance and feedback. We are grateful to Columbia University Environmental Health and Safety Department for providing the chemical inventories. We are thankful to Dr. Ishmail Abdus-Saboor, Dr. Hannah Bayer, Dr. Carol Mason and Dr. Daphna Shohamy for their critical review and insightful comments on this manuscript. We also thank the Zuckerman Institute’s leadership and administration, Dr. Rivka Shoulson for guidance about common drugs used in animal research protocols at Columbia University, as well as laboratory technicians across various labs at the Zuckerman Institute for their participation in this project. Research in this publication was supported by National Institute of Mental Health of the National Institutes of Health under Award Numbers T32-MH018870 and T32-MH126036. The content herein is solely the responsibility of the authors and does not necessarily represent the official views of the National Institutes of Health.

## Author Contributions

EMP, AMZ, DJ conceived the project; EMP, AMZ developed the methodology; AMZ, EMP, CL, DJ, NM, IS, ER, ML performed analysis and literature review; EMP, AMZ wrote the manuscript. All authors reviewed and edited the manuscript.

